# Cerebrovascular Injuries Induce Lymphatic Invasion into Brain Parenchyma to Guide Vascular Regeneration in Zebrafish

**DOI:** 10.1101/346007

**Authors:** Jingying Chen, Jianbo He, Qifen Yang, Yaoguang Zhang, Lingfei Luo

## Abstract

Damage to regional cerebrovascular network and neuronal tissues occurs during acute cerebrovascular diseases, such as ischemic stroke. The promotion of vascular regeneration is the most promising therapeutic approach. To understand cellular and molecular mechanisms underlying brain vascular regeneration, we developed two zebrafish cerebrovascular injury models using genetic ablation and photochemical thrombosis. Although brain parenchyma is physiologically devoid of lymphatic vasculature, we found that cerebrovascular injuries induce rapid ingrowth of meningeal lymphatics into the injured parenchyma. The ingrown lymphatics on one hand become lumenized drain interstitial fluid to resolve brain edema, on the other hand act as “growing tracks” for nascent blood vessels. The ingrown lymphatic vessels undergo apoptosis and clearance after cerebrovascular regeneration. This study reveals a pathological function of meningeal lymphatics, through previously unexpected ingrowth into brain parenchyma and a newly identified lymphatic function as vascular “growing tracks”.

**HIGHLIGHTS:** Cerebrovascular injuries induce lymphatic ingrowth into the injured brain parenchyma The ingrown lymphatics drain interstitial fluid to resolve brain edema Nascent blood vessels use the ingrown lymphatic vessels as “growing tracks” The ingrown lymphatic vessels undergo apoptosis after vascular regeneration completes

## INTRODUCTION

Ischemic stroke, a major cause of death and adult disability, results from brain vascular occlusion and the consequent injury of the regional vascular and neuronal tissues (Krupinski et al., 1993; Liu and Levine, 2008). Promotion of cerebrovascular regeneration, involving the growth of nascent blood vessels (neoangiogenesis) into the post-ischemic infarction area, is a prerequisite to re-nourish the injured brain parenchyma thus becoming the most promising therapeutic approach (Shen et al., 2008; Thored et al., 2007). Angiogenic signals including factors and proteases such as VEGF, PDGF and matrix metalloproteinases have been reported to originate from injured neurons, reactive astrocytes and blood vessels (Greenberg et al., 2013; Su et al., 2008; Heissig et al., 2003). However, studies have yet to be made to understand cellular and molecular mechanisms underlying how the injured brain parenchyma becomes revascularized.

Lymphatic vessels play essential roles in fluid homeostasis, fat absorption, and several pathological processes including lymphedema and tumor metastasis (Alitalo et al., 2005). Lymphatic vessel originate from primitive vein during development (Srinivasan et al., 2007; Nicenboim et al., 2015; Yaniv et al., 2006; Küchler et al., 2006), which is characterized by the drainage of interstitial fluid rather than blood, and the expression of molecular signatures including Prox1, Vegfr3/Flt4, and Lyve1 (Flores et al., 2010; Wigle and Oliver, 1999; Okuda et al., 2012). The lymphangiogenesis is dependent on Vegfc (Karkkainen et al., 2004; Villefranc et al., 2013) and Ccbe1 (Hogan et al., 2009). The latter is required for the processing and maturation of Vegfc (Le Guen et al., 2014; Roukens et al., 2015). Absence of lymphatic vasculature in the central nervous system (CNS) is a long-held concept. Recently, meningeal lymphatics have been discovered in mouse as authentic lymphatic vessels to drain liquid and carry immune cells from the cerebrospinal fluid into the periphery (Aspelund et al., 2015; Louveau et al., 2015). They interact with blood vessels to remove toxic waste (Da Mesquita et al., 2018). In zebrafish, meningeal mural lymphatic ECs (muLECs), also called brain lymphatic endothelial cells (BLECs) or fluorescent granular perithelial (FGP) cells, are able to regulate meningeal angiogensis during development and endocytose macromolecules (Bower et al., 2017; Venero Galanternik et al., 2017; van Lessen et al., 2017). Although muLECs/BLECs/FGPs do not form tubes, they express all lymphatic markers and are sensitive to loss of Ccbe1/Vegfc/Vegfr3, thus believed as zebrafish meningeal lymphatics. The brain parenchyma is devoid of lymphatics under physiological conditions. Whether lymphatic vessels are able to invade brain parenchyma and whether meningeal lymphatics play roles under pathological circumstances are unknown.

The zebrafish model system has substantially contributed to the understanding of cellular and molecular mechanisms underlying organ regeneration based on its powerful regenerative capacity, in particular the organs that are limitedly regenerated in mammals like heart (Jopling et al., 2010; Kikuchi et al., 2010; Zhang et al., 2013). The bacterial nitroreductase (NTR) and its substrate metronidazole (Mtz) have been successfully used to induce conditional targeted cell ablation for regeneration studies in zebrafish (Zhang et al., 2013; Choi et al., 2014; He et al., 2014). However, this system has not been applied to brain vasculature. The photochemically induced thrombosis techniques have been reported to efficiently induce photothrombotic ischemia and damage to local blood vessels in the mouse and rat brain as well as in the zebrafish DA (Maxwell and Dyck, 2005; Kunze et al., 2006; Lee et al., 2007; Kuroiwa et al., 2009; Lee et al., 2017), but have not been applied to zebrafish brain vasculature.

In the present study, we applied the NTR-Mtz system and photochemical thrombosis to introduce injuries to brain vasculature in zebrafish. In response to brain vascular injury, *in vivo* time-lapse imaging revealed that the muLECs/BLECs/FGP cells rapidly invade the injured brain parenchyma, forming lumenized lymphatic vessels. These invaded lymphatic vessels on one hand drain interstitial fluid from the injured area to resolve cerebral edema, on the other hand act as “growing tracks” to provide guidance and support for the growth of nascent blood vessels. The ingrown lymphatic vessels undergo apoptosis after brain vascular regeneration is complete. Hence, the brain parenchyma reverts to the physiological, lymphatic-free status. This temporary ingrown lymphatic vessel-mediated brain vascular regeneration mechanism also occurs in zebrafish photochemical thrombosis model. The results of this study reveal the unexpected ingress of meningeal lymphatics into the brain parenchyma under pathological, vascular injury circumstances. This temporary lymphatic vasculature resolves edema and provides growing tracks for cerebrovascular regeneration.

## RESULTS

### Establishment of a Brain Vascular Injury-Regeneration Model in Zebrafish

To gain insight into cerebrovascular regeneration, we established a zebrafish cerebrovascular injury-regeneration model using the NTR-Mtz system. The *Tg(kdrl:DenNTR)*^*cq10*^ transgenic line was generated, with Dendra2 (Den) fused to NTR and driven by the vascular-specific promoter *kdrl*. Controlled by the *Tg(kdrl:DenNTR)* larvae treated with DMSO (Figure 1A) and *Tg(kdrl:GFP)* larvae treated with Mtz (Figure 1B), incubation of the *Tg(kdrl:DenNTR)* larvae at 3 days post fertilization (3 dpf) with Mtz led to ablation of a majority of regional cerebrovascular endothelial cells (ECs) at 0 days post treatment (0 dpt, equivalent to 4 dpf), but left some larger vessels intact (Figure 1C, arrowheads). Because blood vessel ECs (BECs) in the brain are more sensitive to Mtz than those in the trunk, specific injury to brain blood vessels was achieved using a low concentration (1 mM) of Mtz, without affecting trunk vasculature or generating body phenotypes (Figure S1A). Widespread apoptotic signals were detected in the brain after Mtz treatment, including endothelial and neuronal tissues (Figures S1B and S1C). Local hemorrhages appeared in the brain (Figure S1D and S1E), mimicking the symptoms of massive ischemic stroke. Macrophage infiltration and the engulfment of dead cells were observed (Figures S1F and S1G). Mtz treatment damaged the local cerebrovascular network and generated an injured area in the brain parenchyma. The emergence of neoangiogenesis from the peri-injured into the injured area was observed at 1 dpt (Figure 1C, arrows). This regeneration process proceeded at 2 dpt, became evident at 5 dpt, and completed at 7-8 dpt (Figures 1A–1D). The behaviors, lifespan and fertility of post-regeneration animals showed no difference to the uninjured control, suggesting the brain vascular regeneration as a full functional recovery.

**Figure 1.**
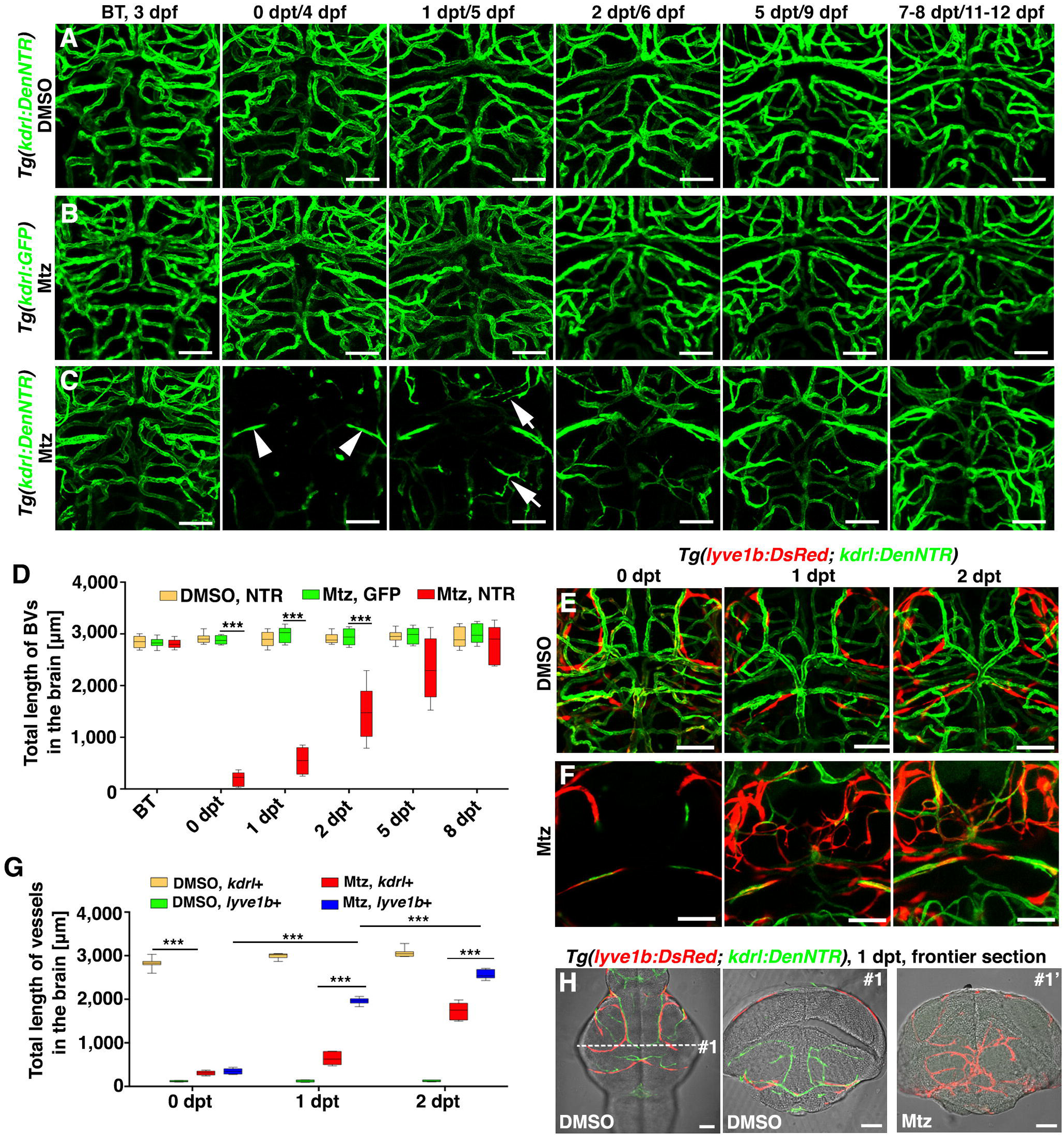
Lymphatic vessels appear in the brain parenchyma after cerebrovascular injury. (A–D) Zebrafish NTR-Mtz cerebrovascular regeneration model. In contrast to the *Tg(kdrl:DenNTR)* larvae treated with DMSO (A, n=40/40) and *Tg(kdrl:GFP)* larvae treated with Mtz (B, n=49/49), brain vasculature of the *Tg(kdrl:DenNTR)* larvae were specifically injured after incubation with 1 mM Mtz (C, 0 dpt, n=39/39). A time course of regeneration is shown. The speed of the neoangiogenesis is presented as the total length of the blood vessels (D, n=10. Two-way ANOVA by Dunnett’s multiple comparisons test. All *P*<0.0001). Arrowheads indicate the remaining relatively large vessels. Arrows indicate the emerging nascent blood vessels. \*\*\**P*<0.001. Data are represented as mean ± SEM. Scale bar, 50 μm. See also Figure S1. (E–G) In the control, the *lyve1b*+ LECs were absent in the brain parenchyma, only present on the optic tectum (E, n=24/24). After Mtz treatment (F, n=20/24), growth of the *lyve1b*+ lymphatics was rapid and active at 1 dpt, but apparently slowed at 2 dpt. The quantifications show the total length of *kdrl*+ and *lyve1b*+ vessels in the brain at 0-2 dpt (G, n=10. Two-way ANOVA by Dunnett’s multiple comparisons test. All *P*<0.0001). \*\*\**P*<0.001. Data are represented as mean ± SEM. Scale bar, 50 μm. See also Figure S2. (H) In frontier sections, the *lyve1*+ lymphatics were predominantly localized in the meninges at the brain surface in the control larvae (#1, n=6/6), whereas became abundant in the brain parenchyma at 1 dpt after Mtz treatment (#1′, n=6/6). Scale bar, 50 μm. See also Figure S2.

### Brain Vascular Injury Induces a Vegfc-Dependent Ingrowth of Meningeal Lymphatics into the Injured Parenchyma

Using the *lyve1b* or *prox1a* promoter-driven transgenes to label lymphatic vessels (Flores et al., 2010; Van Impel et al., 2014), we found rapid and active ingrowth of lymphatics into the injured area on the first day after cerebrovascular injury, but apparently slowed by the second day (Figures 1E–1G, S2, and Video S1). Frontier sections of the brain showed that lymphatics (*lyve1*+) were only present in the meninges at the brain surface under physiological conditions (Figure 1H, #1), but became abundant in the brain parenchyma at 1 dpt after injury (Figure 1H, #1′). Meningeal lymphatics/muLECs turn out to be the only known lymphatic ECs (LECs) inside zebrafish skull (Bower et al., 2017; Venero Galanternik et al., 2017; van Lessen et al., 2017). So, we investigate whether lymphatics in the parenchyma are actually the ingrown meningeal lymphatics after injury. Time-lapse live imaging illustrated a rapid growth of meningeal lymphatics into the injured parenchyma at 0-1 dpt (Figure 2A, and Video S1). The initiation of lymphatic ingrowth and branching at the early phase post-injury was exhibited by 3D-reconstructions (Figure 2B, and Video S2). Cre/loxP-based lineage tracing was performed using the *Tg(lyve1b:CreER*^*T2*^; *β-actin:loxP-RFP-loxP-GFP; lyve1b:DsRed; kdrl:DenNTR)* line (Figure 2C). Negatively controlled by Mtz plus ethanol (Figure 2D) and positively controlled by DMSO plus 4-hydroxytamoxifen (4-OHT; Figure 2E), most ingrown lymphatics were double positive for both GFP and DsRed if Mtz plus 4-OHT were applied (Figures 2F and 2G; STAR Methods). To further substantiate that the ingrown lymphatic vessels emerge from the *lyve1b*+ muLECs, photoconversion of single muLECs loop was fulfilled before Mtz treatment using the *Tg(lyve1b:Kaede*^*cq63*^; *kdrl:CFP-NTR*^*cq62*^) transgenic line. Kaede-red fluorescence was retained in the ingrown vessels (Figures 2H and 2I; STAR Methods), suggesting the photoconverted muLECs as their origins. All the results above demonstrate that meningeal lymphatics respond to cerebrovascular injury and rapidly grow into the injured brain parenchyma.

**Figure 2.**
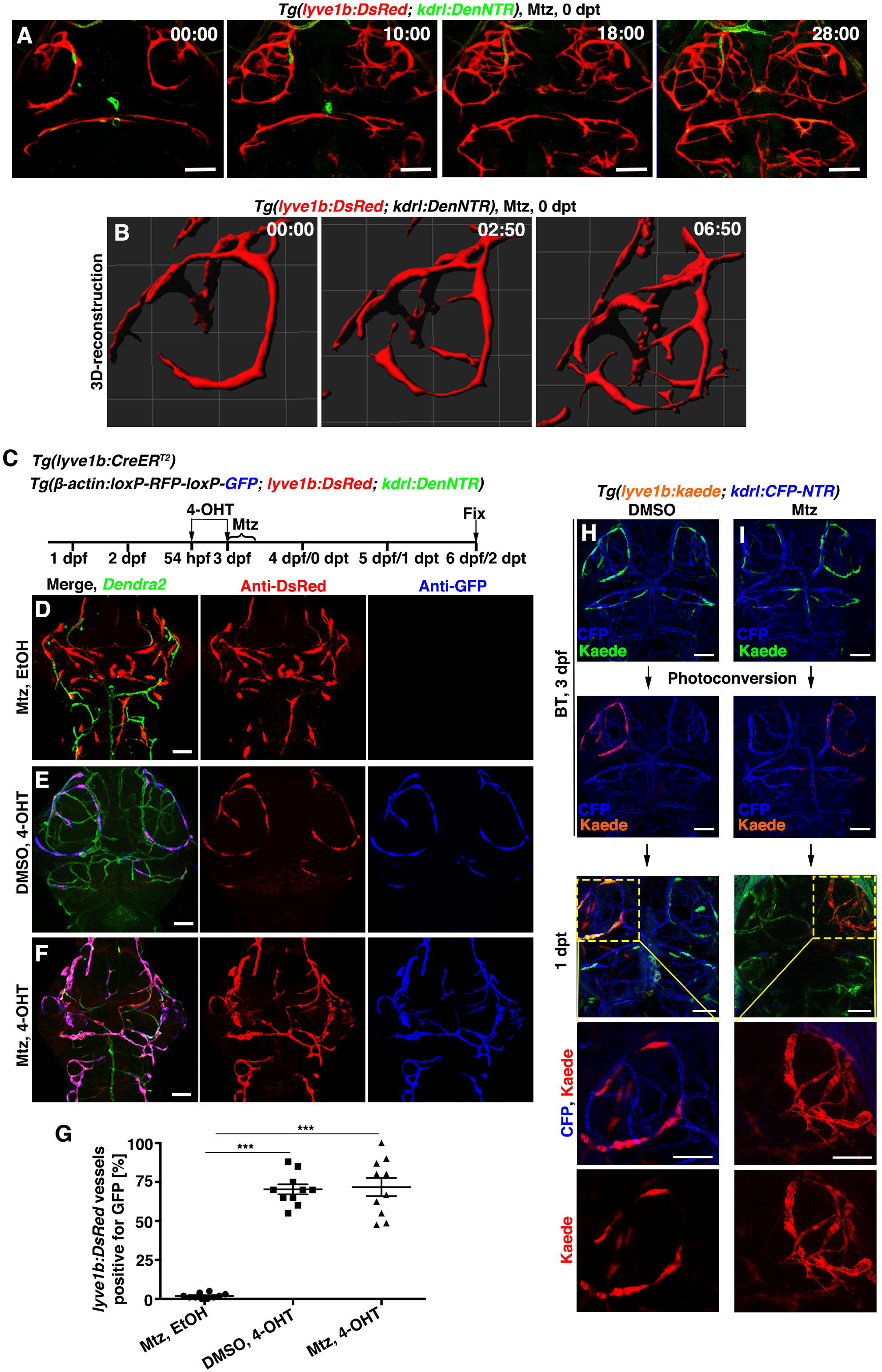
Meningeal lymphatics rapidly grow into the injured parenchyma in response to cerebrovascular damage. (A and B) Time-lapse imaging showed the growth of meningeal lymphatics into the injured area (A, n=10/10). Images at the early phases after Mtz treatment showed that the initiation of ingrowth and branching of meningeal lymphatics by 3D-reconstruction (B, n=10/10). The elapsed time is indicated in hours:minutes after 0 dpt. See also Videos S1 and S2. (C–G) The ingrown vessels are of the lymphatic origin. Transgenic lines, time points of Mtz and 4-OHT administrations were shown in (C). Negatively controlled by Mtz plus ethanol (D, n=12/12) and positively controlled by DMSO plus 4-OHT (E, n=19/21), the ingrown vessels were double positive for anti-DsRed and anti-GFP antibodies if Mtz plus 4-OHT were applied (F, n=21/24). The statistics show the ratios of the *lyve1b:DsRed*-labeled vessels positive for GFP (G, n=10. Two-tailed unpaired t-test, *P*<0.0001). \*\*\**P*<0.001. Data are represented as mean ± SEM. Scale bar, 50 μm. (H and I) In contrast to DMSO treatment (H, n=10/10), photoconversion of muLECs sssfollowed by Mtz treatment resulted in retention of Kaede-red fluorescence in the ingrown vessels (I, n=12/12). BT, Before treatment. Scale bar, 50 μm.

Active lymphangiogenesis requires lymphangiogenic factors. Using *kdrl:DenNTR* and *nkx2.2:GFP* transgenes, fluorescent *in situ* hybridizations (FISH) showed activation of the classical pro-lymphangiogenic marker *vegfc* in the remaining BECs and a portion of *nkx2.2*-positive neurons, inside and at the border of the injured area at 1 dpt (Figures 3A, 3B, and Figures S3A, S3B; STAR Methods). Activation of *vegfc* dramatically reduced at 3 dpt and reached the bottom level at 4-5 dpt (Figure 3C; STAR Methods). Next, we used the *Tg(kdrl:DenNTR; fli1:GFP)* double transgenic line to analyze expression of the Vegfc receptor Vegfr3. The *fli1* promoter is active in both BECs and LECs (Yaniv et al., 2006; Küchler et al., 2006), thus the *fli1*+*kdrl*- vessels are believed to be lymphatic vessels. This Vegfc receptor Vegfr3 was exclusively expressed in the ingrown LECs (*fli1+kdrl-*) and kept absent in the BECs (*kdrl+*) (Figures 3D–3F, and Figures S3C–S3E; STAR Methods). These expression patterns suggest that residual BECs and neurons secrete the pro-lymphangiogenic factor Vegfc to act on LECs, thus promoting the ingrowth of meningeal lymphatics. To make sure the functional significance of Vegfc for the ingrowth of meningeal lymphatics, we applied the *ccbe1* mutant. Ccbe1 is required for the processing and maturation of Vegfc, thereby its mutation causes defective lymphangiogenesis (Hogan et al., 2009; Le Guan et al., 2014; Rouken et al., 2015; van Lessen et al., 2017). Under the *Tg(kdrl:DenNTR; kdrl:mCherry*^*cq15*^; *fli1:GFP)* triple transgenic background, although the *ccbe1* mutant exhibited defects in lymphatic development and slight cerebral edema as indicated by the inter-optic distance, its brain vasculature remained normal (Figures 3G, 3H, 3K, 3L, 3M, 3N, and 3Q). However, ingrowth of meningeal lymphatics (*fli1*+*kdrl*-) and cerebrovascular regeneration (*kdrl*+) failed to occur in the mutant after injury (Figures 3I–3L). The mutant died from severe cerebral edema at 3 dpt (Figures 3O–3Q). These results on one hand ensure the requirement of Vegfc for the meningeal lymphatic ingrowth, on the other hand imply roles of the ingrown lymphatic vessels in edema resolvation and cerebrovascular regeneration. The rest of the study will focus on the functions of the ingrown lymphatics.

**Figure 3.**
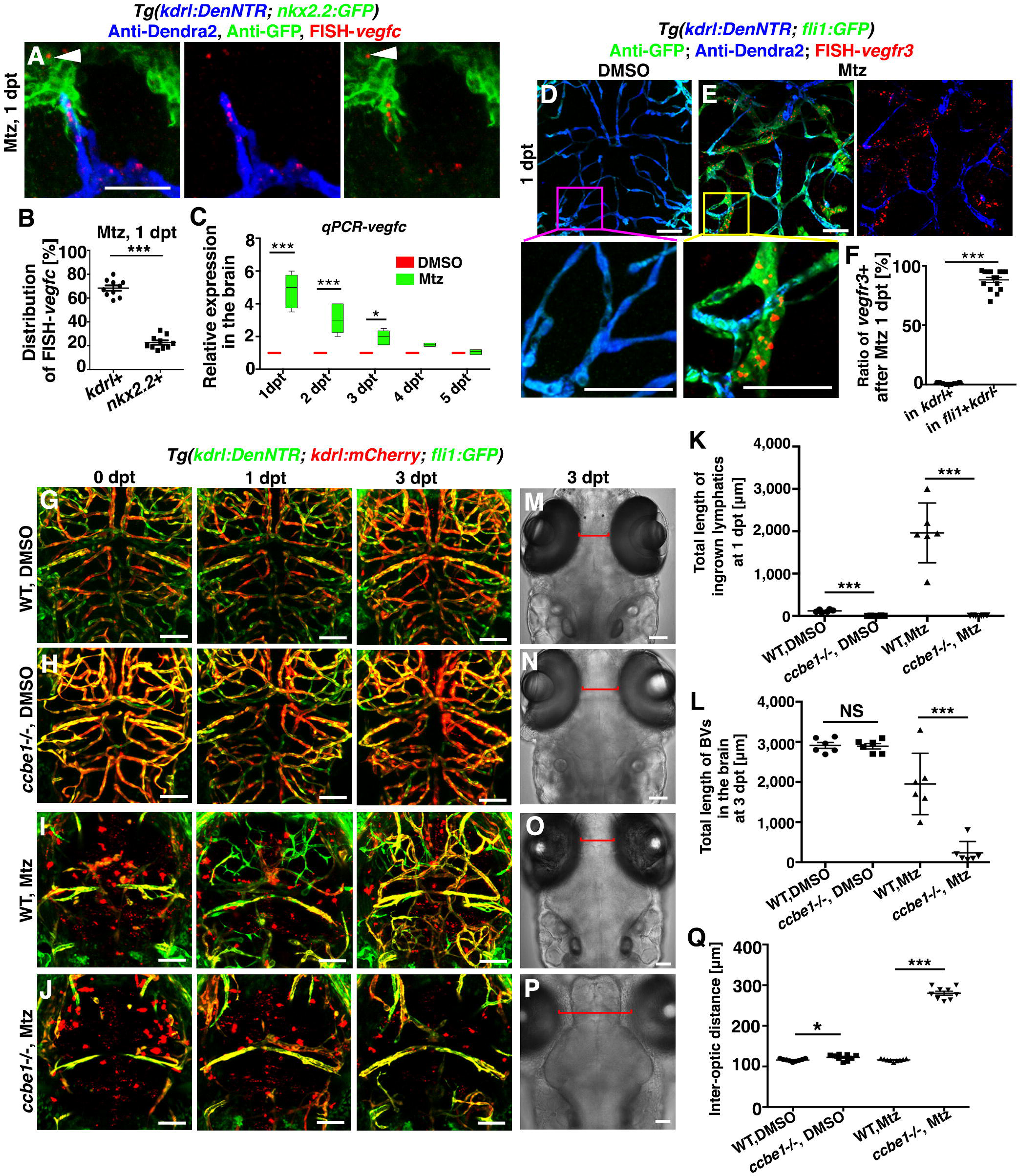
Ingrowth of meningeal lymphatics is dependent on the locally activated Vegfc. (A–C) Local activation of *vegfc*. Expression of *vegfc* at 1 dpt (A, n=30/34). Approximately 68% and 22% of the FISH-*vegfc* signals are distributed in the BECs and *nkx2.2*+ neurons (A, arrowhead), respectively (B, n=10. Two-tailed unpaired *t*-test, *P*<0.0001). RT-qPCRs using dissected brain tissues show the relative expression levels of *vegfc* in the brain at 1-5 dpt (C, n=5. Two-way ANOVA by Dunnett’s multiple comparisons test. DMSO, 1 dpt vs. Mtz, 1 dpt, *P*<0.0001; DMSO, 2 dpt vs. Mtz, 2 dpt, *P*<0.0001; DMSO, 3 dpt vs. Mtz, 3 dpt, *P*=0.0119). Scale bar, 50 μm. See also Figure S3. (D–F) In contrast to the absence of *vegfr3* in the control (D, n=18/20), *vegfr3* is exclusively expressed in the *fli1*+*kdrl*- LECs at 1 dpt (E, n=17/22). Higher magnification images of the framed areas are displayed. The statistics show the ratios of vessels positive for *vegfr3* (F, n=15. Two-tailed unpaired *t*-test, *P*<0.0001). Scale bar, 50 μm. See also Figure S3. (G–L) Brain vascular development was unaffected in the *ccbe1* mutant (H, n=14/14) in contrast to the wild-type (G, n=22/22). After Mtz treatment, meningeal lymphatic ingrowth (*fli1*+*kdrl*-) and neoangiogenesis (*kdrl+*) did not occur in the *ccbe1* mutant (J, n=23/27) in contrast to the wild-type (I, n=36/40). Red spots in (I and J) represents endocytosed kdrl:mCherry red fluorescence by macrophages. The quantifications show the total lengths of the ingrown lymphatics at 1 dpt (K, n=6. Two-tailed unpaired *t*-test, all *P*<0.0001) and nascent blood vessels (BV) at 3 dpt (L, n=6. Two-tailed unpaired *t*-test, WT, DMSO vs. *ccbe1*-/-, DMSO, *P*=0.8383; WT, Mtz vs. *ccbe1*-/-, Mtz, *P*<0.0001). Scale bar, 50 μm. (M–Q) Absence of ingrown lymphatics causes cerebral edema. In contrast to the wild-type (M, n=22/22), the *ccbe1* mutant exhibited slight cerebral edema (N, n=14/14) indicated as the inter-optic distance (red lines). At 3 dpt after Mtz treatment, significantly enlarged inter-optic distance in the *ccbe1* mutant (P, n=8/8) indicated severe cerebral edema in contrast to the wild-type (O, n=8/8), confirmed by statistical analysis (Q, n=9. Two-tailed unpaired *t*-test, WT, DMSO vs. *ccbe1*-/-, DMSO, *P*=0.0209; WT, Mtz vs. *ccbe1*-/-, Mtz, *P*<0.0001). Scale bar, 50 μm. Data are represented as mean ± SEM. \*\*\**P*<0.001, \**P*<0.05, NS, not significant.

### The Ingrown Lymphatics Become Lumenized to Drain Brain Interstitial Fluid

Severe brain vascular injury usually leads to cerebral edema (Greenberg et al., 2013). The lethal edema in the *ccbe1* mutant (Figures 3O–3Q) provides evidences that the ingrown lymphatics drain interstitial fluid from the injured brain to resolve edema. Because zebraifsh meningeal lymphatics are nonlumenized under physiological conditions (Bower et al., 2017; Venero Galanternik et al., 2017; van Lessen et al., 2017), we investigate whether the ingrown lymphatics become lumenized and gain the interstitial fluid drainage function. Controlled by the dorsal aorta (DA) injection of Rhodamine-Dextran that labeled the lumen of blood vessels, intracerebroventricular (ICV) injection labeled brain interstitial fluid. Drainage of Rhodamine was observed in the lumen of *fli1*+*kdrl*- lymphatic vessels immediately after the ICV injection (Figures 4A–4C), demonstrating that lumenized lymphatic vessels have been formed and interstitial fluid drainage function has been obtained at 3 dpt. Then, we performed ICV injection of Alexa647-IgG (150 kDa) at 1 dpt and 2 dpt to learn approximately when the lumenization occurred. Similar to the control (Figure 4D) (van Lessen et al., 2017), the ingrown lymphatics remained nonlumenized and the LECs kept endocytose the Alexa647-IgG macromolecules at 1 dpt (Figure 4F). The endocytosed Alexa647-IgG maintained and accumulated in the LECs thirty minutes after injection (Figures 4E, and 4G). Lumen of the ingrown lymphatics appeared at 2 dpt (Figure 4H, arrows). Alexa647-IgG filled in the lumen immediately after the ICV injection, then became drained out thirty minutes later (Figure 4I). The focused ion beam scanning electron microscopy (FIB-SEM) analyses later will provide further evidences that the ingrown lymphatics were lumenized at 2 dpt (Figure 5H). All these data demonstrate that the ingrown lymphatics undergo a lumenization process between 1 dpt and 2 dpt, thus becoming functional to drain interstitial fluid to resolve cerebral edema.

**Figure 4.**
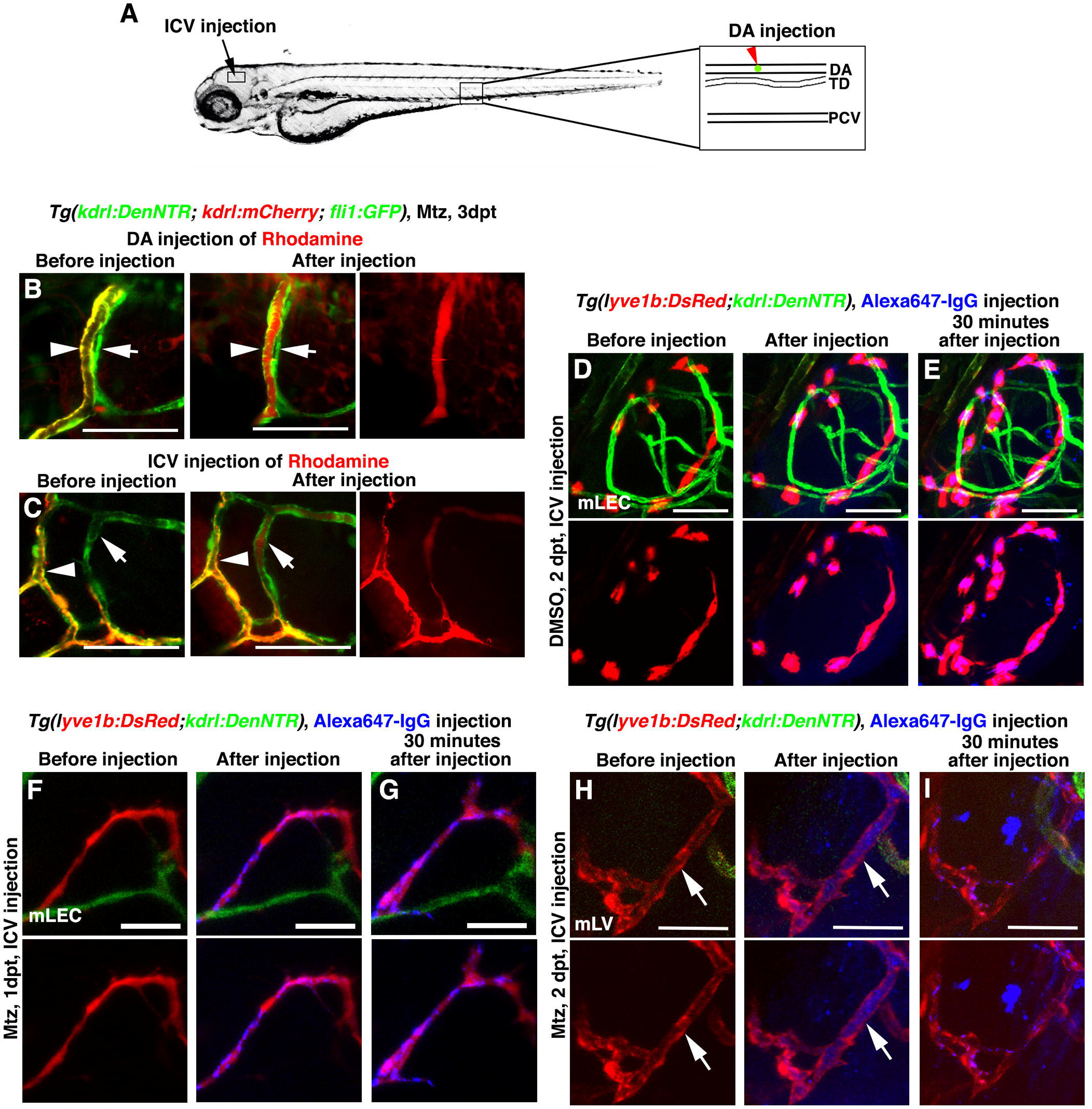
The ingrown lymphatics undergo lumenization to drain brain interstitial fluid. (A-C) The ingrown meningeal lymphatics become lumenized to drain interstitial fluid. Illustrations of DA (dorsal aorta) and ICV (intracerebroventricular) injection points (A). Rhodamine was filled in the blood vessels (arrowheads), but remained absent in the ingrown lymphatics (arrows) after DA injection (B, n=8/10). Drainage of Rhodamine-labeled interstitial fluid was rapidly observed in the ingrown lymphatic vessels (mLVs, arrows) after ICV injection (C, n=15/16). Note that the lumen of mLV was visible (C, arrows). Scale bar, 20 μm. (D–I) Endocytosis, but not lumen flow, of Alexa647-IgG was observed in the meningeal LECs (mLECs) after ICV injection into the control larvae (D, n=15/15), or into the Mtz-treated larvae at 1 dpt (F, n=12/13). Thirty minutes later, the endocytosed dye maintained and accumulated in the mLECs (E, n=14/14; G, n=11/12). If the Alexa647-IgG was injected at 2 dpt after Mtz treatment, the dye filled in the lumen of lymphatic vessels immediately after injection (H, n=17/20). Thirty minutes later, the dye was drained out from the lumen, but the endocytosed dye remained in the mLECs (I, n=16/20). Note that the lumen of lymphatics was clearly visible under confocal microscope at 2 dpt (H, arrows), but not in the control (D) or at 1 dpt (F). Scale bar, 50 μm.

**Figure 5.**
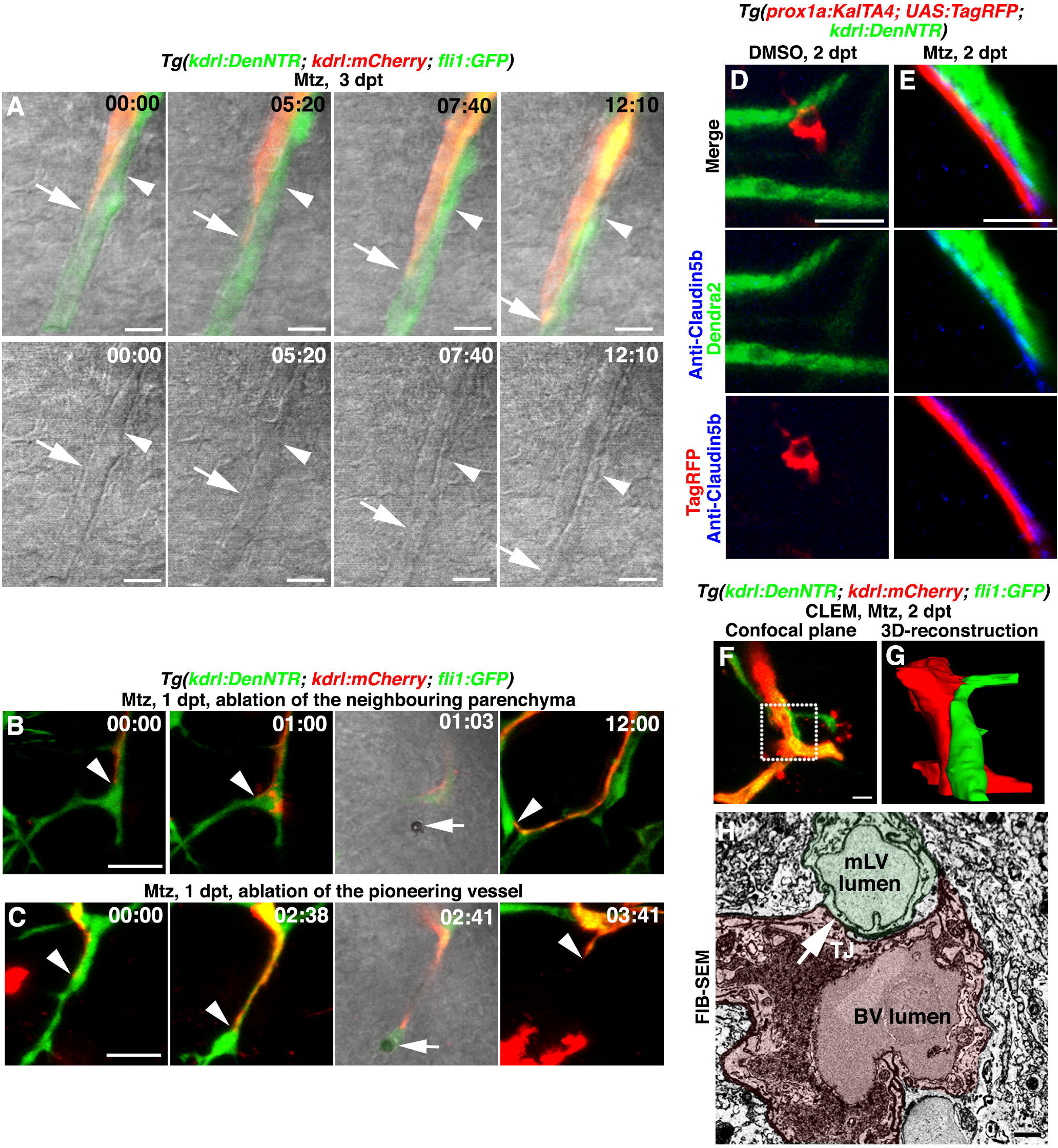
The ingrown lymphatics act as “growing tracks” for nascent blood vessels through direct physical adhesion. (A) Time-lapse imaging showed that the nascent blood vessel (*kdrl*+, arrows) grew along the preformed lymphatic vessel (*fli1*+*kdrl*-, arrowheads). Bright field indicated the two vessel structures. The elapsed time is indicated in hours:minutes. Scale bar, 10 μm. See also Video S3. (B and C) Ablation of the ingrown lymphatic vessel (C, n=20/20 vessels in 10 larvae, 2 vessels per larvae), but not the neighboring parenchyma (B, n=20/20 sites in 10 larvae, 2 sites per larvae), led to the growth arrest and regression of the adhering nascent blood vessel. Arrowheads indicate the laser irradiation sites. The elapsed time is indicated in hours:minutes. Scale bar, 20 μm. See also Video S4. (D and E) Triple labeling of anti-Claudin5b antibodies, TagRFP and Dendra2 epifluosrescences at 2 dpt. In contrast to DMSO treatment (D, n=21/22), Claudin5b was enriched at the interface between the ingrown lymphatic (*prox1a*+) and nascent blood vessels (*kdrl*+) after Mtz treatment (E, n=18/21). Scale bar, 20 μm. See also Figure S4. (F–H) Correlative light and electron microscopy (CLEM) of brain vessels at 6 dpf/2 dpt. The FIB-SEM target areas inside confocal images (F) are indicated with frames. Scale bar, 10 μm. 3D-reconstructions of FIB-SEM single plans identify ingrown meningeal lymphatic vessel (mLV) and nascent blood vessel (BV) as two adhering vessel structures (G, n=3/3). The framed area in (F) is shown in (G). Single FIB-SEM image plane indicates the lumens of mLV and BV (H, n=3/3). Light green and light red mark the cross sections of lymphatic vessel and blood vessel, respectively. Their junctional line represents tight junctions (TJ, arrow). Scale bar, 1 μm. See also Video S5.

### The Ingrown Lymphatics Act as “Growing Tracks” for Nascent Blood Vesels

Based on phenotypes of the *ccbe1* mutant (Figure 3J), we next explore roles of the ingrown lymphatics in brain vascular regeneration. In the *Tg(kdrl:DenNTR; kdrl:mCherry; fli1:GFP)* triple transgenic line, nascent blood vessels (*kdrl+*) were observed growing along the preformed, ingrown lymphatics (*fli1+kdrl-*) using time-lapse imaging (Figure 5A, and Video S3). When the ingrown lymphatics were specifically ablated by the focused irradiation of a high-energy multi-photon laser, the nascent blood vessel growing alongside exhibited growth arrest and regression. In contrast, the ablation of parenchymal tissues neighboring the lymphatics did not affect the nascent blood vessel (Figures 5B, 5C, and Video S4; STAR Methods). These data suggest the ingrown lymphatics as “growing tracks” of nascent blood vessels, most likely through direct physical adhesion. This physical adhesion was supported by the enrichment of surface adhesion molecules Claudin5b and Cdh5 at the interface between the ingrown lymphatic and adjacent blood vessels at 2 dpt (Figures 5D, 5E, and Figure S4), proposing the paracellular interaction between BECs and LECs. Correlative light and electron microscopy (CLEM) and 3D-reconstructions of FIB-SEM substantiated the ingrown lymphatics and nascent blood vessels as two adhering vessel structures with lumens (Figures 5F–5H, and Video S5; STAR Methods). All these results demonstrate that the ingrown lymphatics act as “growing tracks” to support and guide the growth of nascent blood vessels.

### The Ingrown Lymphatic Vessels Undergo Apoptosis and Clearance after Cerebrovascular Regeneration Is Complete

Taken together the data above, meningeal lymphatics rapidly grow into brain parenchyma in response to cerebrovascular damage. On one hand they drain interstitial fluid to resolve brain edema, on the other hand they act as “growing tracks” to guide the growth of nascent blood vessels. However, this ingrown lymphatics-mediated cerebrovascular regeneration mechanism resulted in the presence of lymphatic vessels in the brain parenchyma, inconsistent with its physiologically lymphatic-free status. Therefore, we explored the fate of the ingrown lymphatic vessels after the completion of cerebrovascular regeneration. Time courses using the *lyve1b* or *prox1a* promoter-driven transgenes provided evidences that the ingrown lymphatic vessels gradually disappeared from 5 dpt to 7 dpt (Figure 6A, and S2). Apoptotic signals, detectable in some ingrown LECs (*fli1*+*kdrl*-) from 4 dpt onward (Figure S5), became strong and evident at 6 dpt, but were rarely detectable in the BECs (*kdrl*+) (Figures 6B–6D). These data indicate apoptosis and clearance of the ingrown lymphatic vessels after the completion of cerebrovascular regeneration, so that the brain parenchyma recover the lymphatic-free status.

**Figure 6.**
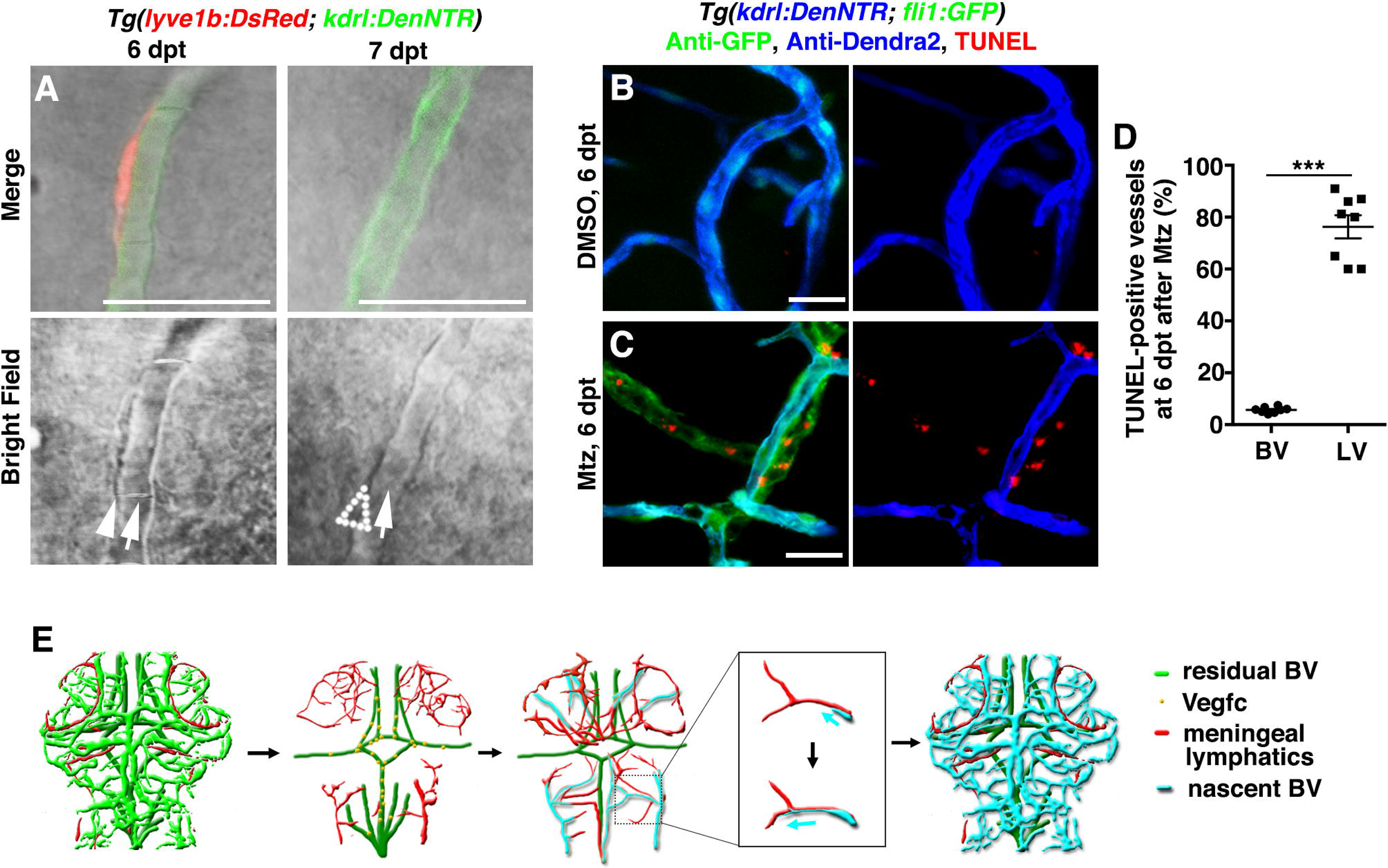
The ingrown lymphatic vessels undergo apoptosis after the completion of cerebrovascular regeneration. (A) The ingrown lymphatic vessels disappeared from 6 dpt to 7 dpt (n=23/26). Arrowhead and arrows indicate the adhering lymphatic and blood vessels, respectively. Note the disappearance of the lymphatic vessel at 7 dpt (dotted arrowhead). Scale bar, 20 μm. (B–D) In contrast to the control (B, n=33/35), TUNEL signals were detected in approximately 70% of the ingrown lymphatic vessels (*fli1*+*kdrl*-) (C, n=30/33), but in less than 5% of the blood vessels (*kdrl*+) (D, n=8. Two-tailed unpaired *t*-test, *P*<0.0001). Scale bar, 20 μm. Data are represented as Mean ± s.e.m., \*\*\**P*<0.001. See also Figure S5. (E) Illustrations of brain vascular regeneration promoted and guided by the temporarily ingrown lymphatics. BV, blood vessel.

### The Lymphatics-Mediated Regeneration Mechanism Is Conserved in Zebrafish Photochemical Thrombosis Model

To answer the question whether the ingrowth of meningeal lymphatics occurs in other cerebrovascular injury models, we established another zebrafish model using photochemical thrombosis, closely mimicking human cerebral thrombosis. Titrations showed that cold light illumination for 600 seconds after injection of Rose bengal induced thrombosis and damage to blood vessels in the illuminated area of zebrafish brain (Figure S6, arrow and arrowhead; Video S6; STAR Methods). In contrast to the control larvae injected with Rhodamine (Figure 7B), meningeal lymphatics (*lyve1*+) invaded the injured area (Figure 7C, dotted circle) by 2 days post illumination (dpi). Nascent blood vessels (*kdrl*+) (Figure 7C, arrowheads) grew along the ingrown lymphatics (Figure 7C, arrow) and cerebrovascular regeneration fulfilled by 4 dpi. From 6 dpi to 7 dpi, the singrown lymphatics were cleared from the injured area and cerebrovascular regeneration accomplished (Figure 7C). These results indicate that the meningeal lymphatics-mediated brain vascular regeneration mechanism is conserved in the photochemical thrombosis model.

**Figure 7.**
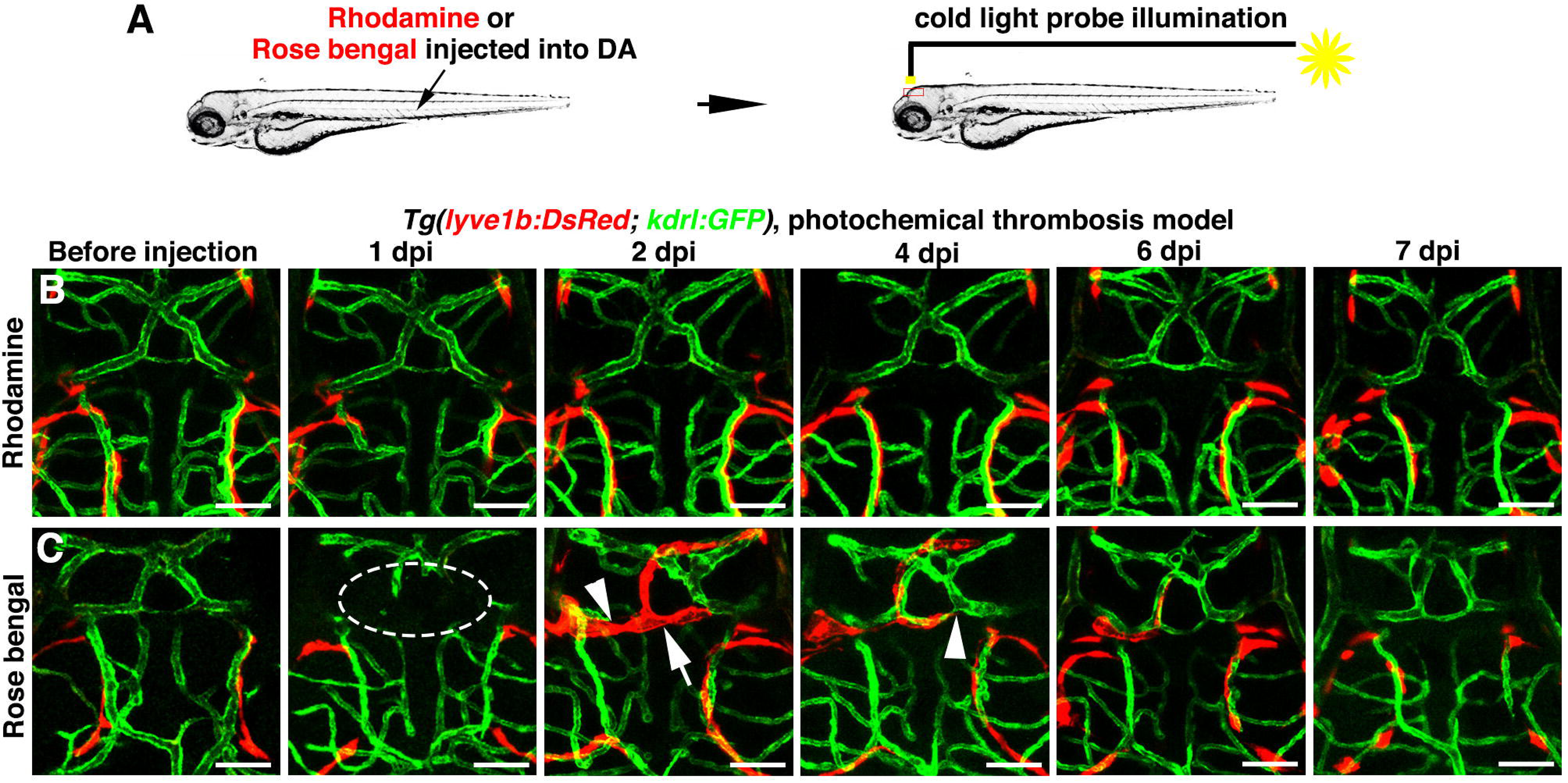
The temporary ingrown lymphatics-mediated cerebrovascular regeneration is conserved in the photochemical brain thrombosis model. (A) Technical procedures of the cold light-induced thrombosis in the zebrafish brain. (B) DA injection of Rhodamine followed by cold light illumination was ineffective to local brain blood vessel (n=10/10). Scale bar, 50 μm. (C) One day post DA injection of Rosa bengal and a 600-second-illumination (1 dpi), local cerebrovascular damage was evident in the illuminated area (dotted circle). By 2 dpi, meningeal lymphatics (arrow) grew into the injured area. Nascent blood vessels grew along the ingrown lymphatics, and vascular regeneration fulfilled by 4 dpi. From 6 dpi to 7 dpi, the ingrown lymphatics become cleared from the injured area (n=15/20). Arrowheads indicate the leading edge of growing blood vessels. Scale bar, 50 μm. See also Figure S6 and Video S6.

## DISCUSSION

In summary, brain vascular injury induces rapid ingrowth of meningeal lymphatics into the injured brain parenchyma, which is dependent on the locally activated Vegfc. These ingrown lymphatics on one hand become lumenized to drain interstitial fluid from the injured parenchyma to resolve cerebral edema. On the other hand, they serve as a migratory scaffold to guide and support the growth of nascent blood vessels (Figure 6E). The meningeal muLEC/BLEC/FGP cells in zebrafish were nonlumenized (Bower et al., 2017; Venero Galanternik et al., 2017; van Lessen et al., 2017). No matter these LECs physiologically belong to true lymphatic system or not, it is reasonable to refer them as meningeal lymphatics in this study because they form authentic, lumenized lymphatic vessels after cerebrovascular injury. Lumenization is not a prerequisite to be acting as “growing tracks”, because growth of nascent blood vessels along nonlumenized lymphatics at 1 dpt and along lumenized lymphatic vessels after 2 dpt have both been observed. Formation of the ingrown lymphatic lumen may be fulfilled through fusion of large vacuoles, which have been reported to be abundant in the muLECs (Bower et al., 2017; van Lessen et al., 2017). Details and mechanisms of lymphatic lumenization need further comprehensive investigations.

In the *ccbe1* mutant, both lymphatic ingrowth and cerebrovascular regeneration were blocked (Figures 3I and 3J). However, we believe that the loss of vascular regeneration is dependent on the loss of lymphatic ingrowth based on the following two reasons. First, although VEGF-C agonizes both VEGFR2 and VEGFR3, which are thought to mediate angiogenesis and lymphangiogenesis, respectively, brain vascular development was unaffected in the *ccbe1* mutant (Figures 3G and 3H). These evidences implicate that the Vegfc signaling is dispensable for the angiogenesis in the brain. Second, when the ingrown lymphatic vessels were ablated by laser, the adhered nascent blood vessels exhibited growth arrest and regression (Figures 5B and 5C), showing dependence of the neoangiogenesis on the ingrown lymphatic vessels.

Although the ingrown lymphatics-mediated brain vascular regeneration mechanism is conserved in the NTR-Mtz and photochemical thrombosis models, we should not conclude that the transient lymphatic vessel is a general response to different types of brain vascular injuries. In a zebrafish cerebrovascular rupture model induced using a focused multi-photon laser, the repair of the rupture is mediated by macrophages through the generation of mechanical traction forces (Liu et al., 2016), but not mediated by the ingrown lymphatics. Two possible reasons might account for this mechanistic difference. First, both the NTR-Mtz and photochemical thrombosis models induce BEC apoptosis in a certain brain area, whereas laser irradiation model induces blood vessel ruptures. These different types of injuries to blood vessels, “apoptosis” vs “rupture”, may explain, at least partially explain, why different vascular repair/regeneration mechanisms are involved. Second, the NTR-Mtz and photochemical models induce severe brain edema inside the closed skull, which may be a critical issue to stimulate lymphatic ingrowth to resolve edema. The laser irradiation model only damages one blood vessel, which does not cause obvious brain edema. So, lymphatic vessels are not required to resolve edema after laser irradiation.

Temporary invasion of meningeal lymphatics prompts a reassessment of the absence of lymphatic vessel in the brain parenchyma, warranting a better understanding of pathological involvements of meningeal lymphatics. “Growing tracks” for nascent blood vessels also expand our knowledge of lymphatic functions. This study in zebrafish provides explanations why VEGF-C and VEGFR3 are up-regulated in the rat photothrombotic stroke model (Gu et al., 2001), and why arteriovenous-malformation human patients express lymphatic-associated genes in the brain (Shoemaker et al., 2014). Development of efficacious angiogenic therapies for ischemic stroke is dependent on a thorough understanding of brain vascular regeneration. This study should be informative for the development of translatable post-ischemic therapies, in particular through the application of lymphangiogenic drugs.

## ACKNOWLEDGMENTS

We thank Markus Affolter, Elke Ober, Ben Hogan, Philip S. Crosier, Li Li, Kenneth S. Zaret, Nathan D. Lawson for fish lines, antibodies, plasmids, and discussions. This work was supported by the National Key Basic Research Program of China (2015CB942800), National Natural Science Foundation of China (91539201, 91739304, 31730060, 31330051), and the 111 Program (B14037).

## AUTHOR CONTRIBUTIONS

L.L., J.C., and Y.Z. designed the experimental strategy, analyzed data, and wrote the manuscript. J.H. performed the combination of FISH and antibody staining. Q.Y. carried out the CLEM. J.C. performed all the other experiments in the study.

## DECLARATION OF INTERESTS

The authors declare no competing interests.

## STAR★METHODS

### CONTACT FOR REAGENT AND RESOURCE SHARING

Further information and requests for reagents may be directed to and will be fulfilled by the Lead Contact, Lingfei Luo.

## EXPERIMENTAL MODEL AND SUBJECT DETAILS

### Animal Strains

Zebrafish strains were raised and maintained under standard laboratory conditions according to Institutional Animal Care and Use Committee protocols. Embryos were treated with 0.003% 1-phenyl-2-thiourea (PTU, Sigma) to inhibit pigment formation. A complete list of the zebrafish strains is provided in the Key Resources Table.

## METHOD DETAILS

### Molecular Cloning

The *kdrl:DenNTR* plasmid was constructed by replacement of the *lfabp* promoter sequence in the *pBluescript-lfabp:DenNTR* plasmid with a 6.5kb *kdrl* promoter cassette. Then, the *kdrl:CFP-NTR* plasmid was constructed by replacement of Dendra2 fragment in the *pBluescript-kdrl:DenNTR* plasmid with CFP sequence. The *kdrl:mCherry* plasmids were constructed by replacement of *Dendra2-NTR* cassette in *pBluescript-kdrl:DenNTR* plasmid with the *mCherry* fragments. The *lyve1b:Kaede* and *lyve1b:CreER*^*T2*^ plasmids were constructed by replacements of the *DsRed* cassette in pTol2-*lyve1:DsRed* plasmid with the *kaede* and *CreER*^*T2*^ fragments, respectively, using the In-fusion technology (TAKARA). The *β-actin:loxP-RFP-loxP-GFP* plasmid was constructed by replacements of the *BFP* and *DsRed* fragments in the *pBS-β-actin:loxP-BFP-loxP-DsRed* plasmid (Addgene, 68972) with the *TagRFP* and *GFP* cassettes, respectively.

### Generation of Transgenic Lines

The *Tg(kdrl:DenNTR)*^*cq10*^*, Tg(kdrl:mCherry)*^*cq15*^, and *Tg(β-actin:loxP-RFP-loxP-GFP)*^*cq39*^, *Tg(kdrl:CFP-NTR)*^*cq62*^ transgenic lines were generated using the pBluescript vector. Constructs flanked by the I-SceI restriction sites were co-injected with I-SceI (NEB) into zebrafish embryos of the AB genetic background at the one-cell stage for transgenesis. The *-5.2lyve1b:DsRed* plasmid (provided by Philip S. Crosier), *lyve1b:Kaede* plasimd, and *lyve1b:CreER*^*T2*^ plasmid were co-injected with the capped *Tol2 transposase* RNA (40-50 pg) for transgenesis (*cq27*, *cq63* and *cq25*).

### *In Situ* Hybridizations, Antibody Staining, and Imaging

Whole-mount *in situ* hybridizations were carried out as previously described (Liu et al., 2009) using the *vegfc*, *vegfr3* probes (Figures S2B, S2D). Images were captured using a SteREO DiscoveryV20 microscope equipped with AxioVision Rel 4.8.2 software (Carl Zeiss).

Zebrafish whole-mount antibody staining was performed as previously described (Lu et al., 2013). Primary antibodies against Claudin5b (1:200, Invitrogen), GFP (1:500, Abcam, Santa Cruz), Dendra2 (1:500, Antibody-online), zfCdh5 (Blum et al., 2008) (1:250, a gift from Markus Affolter), mCherry (1:500, Abcam), DsRed (1:500, Aviva Systems Biology) were applied. Secondary antibodies used in the study included donkey anti-goat IgG Alexa fluor 488-conjugated (1:1000, Invitrogen), donkey anti-goat IgG Alexa fluor 633-conjugated (1:1000, Invitrogen), donkey anti-mouse IgG Alexa fluor 568-conjugated (1:1000, Invitrogen), donkey anti-rabbit IgG Alexa fluor 647-conjugated (1:1000, Invitrogen), donkey anti-rabbit IgG Alexa fluor 488-conjugated (1:1000, Invitrogen), donkey anti-mouse IgG Alexa fluor 647-conjugated (1:1000, Invitrogen), goat anti-rabbit IgG Alexa fluor 405-conjugated (1:1000, Invitrogen). Antibody stained or live larvae were mounted in 1.2% low melting point agarose and imaged using ZEN2010 software equipped on an LSM780 confocal microscope (Carl Zeiss).

### Chemical Treatment

For Mtz treatment, larvae at 3 dpf (days post fertilization) were incubated with 1 mM Mtz (Sigma) in 0.2% DMSO for five hours, washed three times with egg water, and recovered in the egg water with 0.003% PTU. Nineteen hours after withdrawal of Mtz was defined as 0 dpt, which is equivalent to 4 dpf.

4-Hydroxytamoxifen (4-OHT, Sigma) was dissolved in 100% ethanol to prepare a stock concentration of 10 mM. Larvae were incubated in 5 μM working solution at the indicated time frame using 0.05% ethanol in the egg water as control (Figures 2C-2F).

### Laser Microsurgical Cell Ablation, Photoconversion and *In Vivo* Time-Lapse Imaging

For laser microsurgical cell ablation (Figures 5B, 5C, and Video S4), the larvae were mounted in 1.2% low melting point agarose in the egg water with 0.003% PTU. The targeted endothelial cell or parenchymal tissue was focused in a single confocal plane using a LSM780NLO multi-photon microscope (Carl Zeiss), Multi-photon laser at 800 nm was generated using a Chameleon Vision 2 Laser System (Coherent) to irradiate the focusing site for 3 seconds.

For photoconversion (Figures 2H and 2I), the *Tg(lyve1b:Kaede; kdrl:CFP-NTR)* larvae were mounted in 1.2% low melting point agarose. The focused muLECs loop regions were irradiated for 30 seconds by the 405-nm laser. The Kaede epifluorescence was converted from green to red after photoconversion.

For time-lapse live imaging, zebrafish embryos were grown in the presence of 0.003% PTU to avoid pigmentation. Larvae were mounted in 1.2% low melting point agarose in the egg water with 0.003% PTU using 35-mm glass bottom dishes. Time-lapse images were captured using a 20× water immersion objective mounted on the LSM780NLO confocal microscope equipped with a heating stage to maintain 28.5 °C. Z image stacks were collected every 5-10 minutes, and three-dimensional data sets were compiled using ZEN2010 software (Carl Zeiss). Selected data sets were 3D-reconstructed and segmented by the iMaris software (Bitplane).

### Combination of Fluorescent *In Situ* Hybridization (FISH) and Antibody Staining

The combination of FISH and antibody staining was performed as previously described (He et al., 2014). In short, larvae were fixed in 4% paraformaldehyde. Then, the skins were manually removed and the larvae were dehydrated in 100% methanol at -20 °C for at least 24 hours. After treatment of 3% H_2_O_2_ in methanol, larvae were serially transferred into 75%, 50%, 25%, and 0% methanol in PBST (1% Triton X-100 in PBS). Next, the larvae were pre-hybridized in the HYB buffer (50% formamide, 5×SSC, 0.1% Tween-20, 5 mg/ml torula yeast RNA, 50 mg/ml heparin) and hybridized with the digoxigenin-labeled *vegfr3*, *vegfc* probes at 63 °C or 65 °C overnight. After removal of probes, the larvae were serially washed with SSCT and MABT (150 mM maleic acid, 100 mM NaCl, 0.1% Tween-20, pH 7.5), then blocked in 2% Block Reagent (Roche), and incubated with the Anti-digoxigenin POD antibodies (1:500, Roche) overnight. The larvae were serially washed MABT, PBST, and PBS, then incubated in TSA Plus Cy5 Solution (Perkin Elmer) overnight and washed again with PBST. Afterwards, antibody staining proceeded as described above. The *Tg(kdrl:DenNTR; fli1:GFP)* and *Tg(kdrl:DenNTR; nkx2.2:GFP)* lines were subjected to antibody staining using the anti-GFP (1:1000, Santa Cruz, #sc9996) and anti-Dendra2 (1:1000, Antibody-online, #ABIN361314) primary antibodies. Then, the goat anti-mouse IgG Alexa fluor 633-conjugated (1:1000, Invitrogen, #A21052) and goat anti-rabbit IgG Alexa fluor 405-conjugated (1:1000, Invitrogen, #A31556) secondary antibodies were used to label GFP and Dendra2, respectively (Figures 3A, 3D, 3E).

### Quantitative Real-time RCR

The total RNAs were extracted from whole brains dissected from one hundred larvae for *vegfc* analysis. Total RNAs were prepared using the TRIzol reagent (Roche). cDNAs were synthesized using the Reverse Transcription Kit (Promega). Quantitative real-time PCRs (qPCRs) were carried out for *vegfc* using the FastStart Universal SYBR Green Master (Roche), normalized by transcriptions of *β-actin* (Figure 3C). A complete list of the primers used in this study is provided in the Key Resources Table.

### Correlative Light and Electron Microscopy (CLEM) and 3D-Reconstructions

The fixed larvae were embedded in the 4% low melting agarose and subjected to 50 μm sections using VT1000S vibratome (Leica). Then, the sections were pasted in a photo-etched grid-50 μ-dish (ibidi) and the lymphatic and vascular ECs with their location in relation to the grid were identified under an LSM780 confocal microscope equipped with ZEN2010 software (Carl Zeiss). In the grid-dish, the sections were fixed in 2.5% glutaraldehyde, 2% formaldehye in Cacodylate buffer (0.1 M cacodylate in water, pH7.0) at 4 °C overnight, washed with Cacodylate buffer for several times, incubated in osmium tetroxide for several times, and stained with 1% uranyl acetate and lead aspartate.

Afterwards, the samples were dehydrated and embedded in Epon resin. The embedded sample was mounted on the SEM sample stub with colloidal graphite, tilted with 54 degree in the chamber and moved to the location in relation to the grid. 3D dataset was acquired by FIB (Focused Ion Beam)-SEM Crossbeam 540 (Carl Zeiss). FIB milled the sample surface in perpendicular direction. FIB worked at 30 kV and firstly at beam current of 15∼30 pA for coarse milling, then at 3 nA for fine milling. Afterwards, FIB worked for continuously serial section at 3 nA, simultaneously the electron beam imaged the latest milled section at 2 kV, 5000 magnification, and beam current of 500 pA. The acquired EM dataset had 25 nm pixel resolution in X/Y scale and 100 nm in Z scale, which was determined by milling interval of FIB. The 3D-reconstruction dataset was calibrated, reconstructed and segmented by ORS Visual software (Carl Zeiss) (Figures 5H and 5G). Movies were rendered by ORS Visual (Video S5).

### Dye Injection and Microangiography

The dye injection and microangiography were performed as previously described (Yaniv et al., 2006; Küchler et al., 2006; Bower et al., 2017; Venero Galanternik et al., 2017; van Lessen et al., 2017). In brief, for angiography, a suspension of IgG-conjugated Alexa Fluor 647 (2 mg/ml, A31573, Invitrogen) or Rhodamine-Dextran 10 kDa (2 mg/ml, D1824, Invitrogen) was injected directly into the dorsal aorta, or intracerebroventricle using glass capillary needles.

### TUNEL Assay

Larvae were fixed in 4% paraformaldehyde in PBS at 4 °C overnight, subjected to skin removal, and assayed using the *In Situ* Cell Death Detection Kit, TMR Red (Roche) according to the manufacturer’s instruction.

### Zebrafish Cerebrovascular Injury by Photochemical Thrombosis

Larvae at 5 dpf were anesthetized with 0.016% ethyl 3-aminobenzoate methanesulfonate (MS-222) and mounted in 1.2% low melting point agarose, followed by the injection of Rose Bengal (150 μg/ml, 632-69-9, BBI) or Rhodamine-Dextran into the dorsal aorta as previously described (Lee et al., 2017). Then, the larvae head was exposed to a self-made light probe connecting to a cold light source (Zeiss, CL6000, LED) for 300 seconds or 600 seconds as previously described (Yu and Li, 2016) (Figures 7 and S6).

## QUANTIFICATION AND STATISTICAL ANALYSIS

All statistical calculations were performed using Graphpad Prism. Variance for all groups data is presented as ± s.e.m. Larvae were collected from incrosses and outcrosses of several pairs of adult zebrafish. Mutant and sibling larvae were grown together in a single tank. Phenotyping preceded genotyping in mutant analyses, hence analysis was genotype blinded. In the other experiments, the investigators were not blinded to group allocation during data collection and/or analysis. All experiments comparing treatment groups were carried out using randomly assigned siblings. After at least two repeated experiments, data were analysed for statistical significance using Two-way ANOVA by Dunnett’s multiple comparisons test and two-tailed upaired t-test. A value of *P*<0.05 was considered to be statistically significant. No data were excluded from analyses. The exact sample size (n), *P* value for each experimental group and statistical tests were indicated in the figure legends.

## Supplemental Video Legends

**Video S1. Meningeal lymphatics rapidly grow into the injured area in response to cerebrovascular damage, Related to Figure 2.** After cerebrovascular injury, the meningeal lymphatics labeled in red speedily responded to initiate growth and branching. Their ingrowth rapidly invaded the injured brain area. The *Tg(lyve1b:DsRed; kdrl:DenNTR)* transgenic line at 0-1 dpt was used for the time-lapse image (n=3/3).Duration of imaging was 2115 minutes. Scale bar, 50 μm.

**Video S2. Early phases of meningeal lymphatic ingrowth, Related to Figure 2.** 3D-reconstructions of meningeal lymphatics at the early phases after Mtz treatment (0 hpt, 2.8 hpt, and 6.8 hpt) validated the ingrowth of meningeal lymphatics/muLECs into the injured area. The *Tg(lyve1b:DsRed; kdrl:DenNTR)* transgenic line was used for the imaging. hpt, hours post treatment.

**Video S3. Nascent blood vessels use the preformed ingrown lymphatics as “growing tracks”, Related to Figure 5.** The nascent blood vessel fluorescently labeled in red grew along the preformed, ingrown lymphatic vessel fluorescently labeled in green. Arrows indicate the leading edge of the growing blood vessel. The *Tg(kdrl:DenNTR; kdrl:mCherry; fli1:GFP)* transgenic line at 3 dpt was used for the time-lapse image (n=15/17). Durations of imaging were 120 minutes. Scale bar, 20 μm.

**Video S4. Ablation of the ingrown lymphatics leads to growth arrest and regression of the adhering blood vessel, Related to Figure 5.** The ablation of parenchymal tissues neighbouring the ingrown lymphatics by focused high energy multi-photon laser was ineffective to the neoangiogenesis (left side, n=20/20 sites). In contrast, if the lymphatic vessel was specifically ablated, the adhering nascent blood vessel showed growth arrest and regression (right side, n=20/20 vessels). Dotted circles and arrows indicate the laser irradiation sites. Arrowheads indicate the leading edge of nascent blood vessels. The *Tg(kdrl:DenNTR; kdrl:mCherry; fli1:GFP)* transgenic line at 1 dpt was used for the time-lapse image. Durations of left and right imaging were 720 minutes and 228 minutes, respectively. Scale bar, 50 μm.

**Video S5. 3D-reconstructions of focused ion beam scanning electron microscopy (FIB-SEM) images, Related to Figure 5.** The acquired SEM datasets had 25 nm pixel resolution in X/Y scale and 100 nm in Z scale. A total of 213 single plan images were used for the 3D-reconstructions of the ingrown lymphatic vessel (green) and the adhering nascent blood vessel (red) in the brain (n=3/3).

**Video S6. Photochemically induced thrombosis in the zebrafish brain, Related to Figure 7 and Figure S6**. The left side sample was injected with Rose bengal, while the right side control was injected with Rhodamine. Erythrocytes were circulated before injection. Blood was labeled by red fluorescence immediately after injection. When the imaged region was illuminated by cold light for 600 seconds, erythrocytes were clogged and thrombosis was induced in the illuminated area. Note that the local blood vessels became damaged. The *Tg(kdrl:GFP; gata1:DsRed)* transgenic line was used for the time-lapse image. Durations of both left and right imaging were 1290 seconds.

## REFERENCES

Alitalo, K., Tammela, T., and Petrova, T.V. (2005). Lymphangiogenesis in development and human disease. Nature 438, 946–953.

Aspelund, A., Antila, S., Proulx, S.T., Karlsen, T.V., Karaman, S., Detmar, M., Wiig, H., and Alitalo, K. (2015). A dural lymphatic vascular system that drains brain interstitial fluid and macromolecules. J. Exp. Med. 212, 991–999.

Beis, D., Bartman, T., Jin, S.W., Scott, I.C., D’Amico, L.A., Ober, E.A., Verkade, H., Frantsve, J., Field, H.A., Wehman, A., et al. (2005). Genetic and cellular analyses of zebrafish atrioventricular cushion and valve development. Development 132, 4193–4204.

Blum, Y., Belting, H.G., Ellertsdottir, E., Herwig, L., Luders, F., and Affolter, M. (2008). Complex cell rearrangements during intersegmental vessel sprouting and vessel fusion in the zebrafish embryo. Dev. Biol. 316, 312–322.

Bower, N.I., Koltowska, K., Pichol-Thievend, C., Virshup, I., Paterson, S., Lagendijk, A.K., Wang, W., Lindsey, B.W., Bent, S.J., Baek, S., et al. (2017). Mural lymphatic endothelial cells regulate meningeal angiogenesis in the zebrafish. Nat. Neurosci. 20, 774–783.

Busch-Nentwich, E., Kettleborough, R., Harvey, S., Collins, J., Ding, M., Dooley, C., Fenyes, F., Gibbons, R., Herd, C., Mehroke, S., et al. (2012). Sanger institute zebrafish mutation project mutant, phenotype and image data submission. ZFIN direct data Submission (http://zfin.org).

Choi, T.Y., Ninov, N., Stainier, D.Y., and Shin, D. (2014). Extensive conversion of hepatic biliary epithelial cells to hepatocytes after near total loss of hepatocytes in zebrafish. Gastroenterology 146, 776–788.

Da Mesquita, S., Louveau, A., Vaccari, A., Smirnov, I., Cornelison, R.C., Kingsmore, K.M., Contarino, C., Onengut-Gumuscu, S., Farber, E., Raper, D., et al. (2018). Functional aspects of meningeal lymphatics in ageing and Alzheimer’s disease. Nature 560, 185–191.

Flores, M.V., Hall, C.J., Crosier, K.E., and Crosier, P.S. (2010). Visualization of embryonic lymphangiogenesis advances the use of the zebrafish model for research in cancer and lymphatic pathologies. Dev. Dyn. 239, 2128–2135.

Greenberg, D.A., and Jin, K. (2013). Vascular endothelial growth factors (VEGFs) and stroke. Cell. Mol. Life Sci. 70, 1753–1761.

Gu, W., Brannstrom, T., Jiang, W., Bergh, A., and Wester, P. (2001). Vascular endothelial growth factor-A and -C protein up-regulation and early angiogenesis in a rat photothrombotic ring stroke model with spontaneous reperfusion. Acta. Neuropathol. 102, 216–226.

He, J., Lu, H., Zou, Q., and Luo, L. (2014). Regeneration of liver after extreme hepatocyte loss occurs mainly via biliary transdifferentiation in zebrafish. Gastroenterology 146, 789–800.

Heissig, B., Hattori, K., Friedrich, M., Rafii, S., and Werb, Z. (2003). Angiogenesis: vascular remodeling of the extracellular matrix involves metalloproteinases. Curr. Opin. Hematol. 10, 136–141.

Hogan, B.M., Bos, F.L., Bussmann, J., Witte, M., Chi, N.C., Duckers, H.J., and Schulte-Merker, S. (2009). Ccbe1 is required for embryonic lymphangiogenesis and venous sprouting. Nat. Genet. 41, 396–398.

Jopling, C., Sleep, E., Raya, M., Marti, M., Raya, A., and Izpisua Belmonte, J.C. (2010). Zebrafish heart regeneration occurs by cardiomyocyte dedifferentiation and proliferation. Nature 464, 606–609.

Karkkainen, M.J., Haiko, P., Sainio, K., Partanen, J., Taipale, J., Petrova, T.V., Jeltsch, M., Jackson, D.G., Talikka, M., Rauvala, H., et al. (2004). Vascular endothelial growth factor C is required for sprouting of the first lymphatic vessels from embryonic veins. Nat. Immunol. 5, 74–80.

Kikuchi, K., Holdway, J.E., Werdich, A.A., Anderson, R.M., Fang, Y., Egnaczyk, G.F., Evans, T., Macrae, C.A., Stainier, D.Y., and Poss, K.D. (2010). Primary contribution to zebrafish heart regeneration by gata4(+) cardiomyocytes. Nature 464, 601–605.

Krupinski, J., Kaluza, J., Kumar, P., Wang, M., and Kumar, S. (1993). Prognostic value of blood vessel density in ischaemic stroke. Lancet 342, 742.

Küchler, A.M., Gjini, E., Peterson-Maduro, J., Cancilla, B., Wolburg, H., and Schulte-Merker, S. (2006). Development of the zebrafish lymphatic system requires VEGFC signaling. Curr. Biol. 16, 1244–1248.

Kunze, A., Grass, S., Witte, O.W., Yamaguchi, M., Kempermann, G., and Redecker, C. (2006). Proliferative response of distinct hippocampal progenitor cell populations after cortical infarcts in the adult brain. Neurobiol. Dis. 21, 324–332.

Kuroiwa, T., Xi, G., Hua, Y., Nagaraja, T.N., Fenstermacher, J.D., and Keep, R.F. (2009). Development of a rat model of photothrombotic ischemia and infarction within the caudoputamen. Stroke 40, 248–253.

Le Guen, L., Karpanen, T., Schulte, D., Harris, N.C., Koltowska, K., Roukens, G., Bower, N.I., van Impel, A., Stacker, S.A., Achen, M.G., et al. (2014). Ccbe1 regulates Vegfc-mediated induction of Vegfr3 signaling during embryonic lymphangiogenesis. Development 141, 1239–1249.

Lee, I.J., Yang, Y.C., Hsu, J.W., Chang, W.T., Chuang, Y.J., and Liau, I. (2017). Zebrafish model of photochemical thrombosis for translational research and thrombolytic screening in vivo. J. Biophotonics 10, 494–502.

Lee, J.K., Park, M.S., Kim, Y.S., Moon, K.S., Joo, S.P., Kim, T.S., Kim, J.H., and Kim, S.H. (2007). Photochemically induced cerebral ischemia in a mouse model. Surg. Neurol. 67, 620–625.

Liu, C., Wu, C., Yang, Q., Gao, J., Li, L., Yang, D., and Luo, L. (2016). Macrophages Mediate the Repair of Brain Vascular Rupture through Direct Physical Adhesion and Mechanical Traction. Immunity 44, 1162–1176.

Liu, S., and Levine, S.R. (2008). The Continued Promise of Neuroprotection for Acute Stroke Treatment. J. Exp. Stroke Transl. Med. 1, 1–8.

Liu, X., Huang, S., Ma, J., Li, C., Zhang, Y., and Luo, L. (2009). NF-kappaB and Snail1a coordinate the cell cycle with gastrulation. J. Cell Biol. 184, 805–815.

Louveau, A., Smirnov, I., Keyes, T.J., Eccles, J.D., Rouhani, S.J., Peske, J.D., Derecki, N.C., Castle, D., Mandell, J.W., Lee, K.S., et al. (2015). Structural and functional features of central nervous system lymphatic vessels. Nature 523, 337–341.

Lu, H., Ma, J., Yang, Y., Shi, W., and Luo, L. (2013). EpCAM is an endoderm-specific Wnt derepressor that licenses hepatic development. Dev. Cell 24, 543–553.

Maxwell, K.A., and Dyck, R.H. (2005). Induction of reproducible focal ischemic lesions in neonatal mice by photothrombosis. Dev. Neurosci. 27, 121–126.

Nicenboim, J., Malkinson, G., Lupo, T., Asaf, L., Sela, Y., Mayseless, O., Gibbs-Bar, L., Senderovich, N., Hashimshony, T., Shin, M., et al. (2015). Lymphatic vessels arise from specialized angioblasts within a venous niche. Nature 522, 56–61.

Okuda, K.S., Astin, J.W., Misa, J.P., Flores, M.V., Crosier, K.E., and Crosier, P.S. (2012). lyve1 expression reveals novel lymphatic vessels and new mechanisms for lymphatic vessel development in zebrafish. Development 139, 2381–2391.

Roman, B.L., Pham, V.N., Lawson, N.D., Kulik, M., Childs, S., Lekven, A.C., Garrity, D.M., Moon, R.T., Fishman, M.C., Lechleider, R.J., et al. (2002). Disruption of acvrl1 increases endothelial cell number in zebrafish cranial vessels. Development 129, 3009–3019.

Roukens, M.G., Peterson-Maduro, J., Padberg, Y., Jeltsch, M., Leppanen, V.M., Bos, F.L., Alitalo, K., Schulte-Merker, S., and Schulte, D. (2015). Functional Dissection of the CCBE1 Protein: A Crucial Requirement for the Collagen Repeat Domain. Circ. Res. 116, 1660–1669.

Shen, F., Fan, Y., Su, H., Zhu, Y., Chen, Y., Liu, W., Young, W.L., and Yang, G.Y. (2008). Adeno-associated viral vector-mediated hypoxia-regulated VEGF gene transfer promotes angiogenesis following focal cerebral ischemia in mice. Gene Ther. 15, 30–39.

Shoemaker, L.D. et al (2014). Human brain arteriovenous malformations express lymphatic-associated genes. Ann. Clin. Transl. Neurol. 1, 982–995.

Srinivasan, R.S., Dillard, M.E., Lagutin, O.V., Lin, F.-J., Tsai, S., Tsai, M.-J., Samokhvalov, I.M., and Oliver, G. (2007). Lineage tracing demonstrates the venous origin of the mammalian lymphatic vasculature. Genes Dev. 21, 2422–2432.

Su, E.J., Fredriksson, L., Geyer, M., Folestad, E., Cale, J., Andrae, J., Gao, Y., Pietras, K., Mann, K., Yepes, M., et al. (2008). Activation of PDGF-CC by tissue plasminogen activator impairs blood-brain barrier integrity during ischemic stroke. Nat. Med. 14, 731–737.

Thored, P., Wood, J., Arvidsson, A., Cammenga, J., Kokaia, Z., and Lindvall, O. (2007). Long-term neuroblast migration along blood vessels in an area with transient angiogenesis and increased vascularization after stroke. Stroke 38, 3032–3039.

van Lessen, M., Shibata-Germanos, S., van Impel, A., Hawkins, T.A., Rihel, J., and Schulte-Merker, S. (2017). Intracellular uptake of macromolecules by brain lymphatic endothelial cells during zebrafish embryonic development. eLife 6.

van Impel, A., Zhao, Z., Hermkens, D.M., Roukens, M.G., Fischer, J.C., Peterson-Maduro, J., Duckers, H., Ober, E.A., Ingham, P.W., and Schulte-Merker, S. (2014). Divergence of zebrafish and mouse lymphatic cell fate specification pathways. Development 141, 1228–1238.

Venero Galanternik, M., Castranova, D., Gore, A.V., Blewett, N.H., Jung, H.M., Stratman, A.N., Kirby, M.R., Iben, J., Miller, M.F., Kawakami, K., et al. (2017). A novel perivascular cell population in the zebrafish brain. eLife 6.

Villefranc, J.A., Nicoli, S., Bentley, K., Jeltsch, M., Zarkada, G., Moore, J.C., Gerhardt, H., Alitalo, K., and Lawson, N.D. (2013). A truncation allele in vascular endothelial growth factor c reveals distinct modes of signaling during lymphatic and vascular development. Development 140, 1497–1506.

Wigle, J.T., and Oliver, G. (1999). Prox1 function is required for the development of the murine lymphatic system. Cell 98, 769–778.

Yaniv, K., Isogai, S., Castranova, D., Dye, L., Hitomi, J., and Weinstein, B.M. (2006). Live imaging of lymphatic development in the zebrafish. Nat. Med. 12, 711–716.

Yu, X., and Li, Y.V. (2016). Zebrafish (Danio rerio) developed as an alternative animal model for focal ischemic stroke. Acta Neurochir. Suppl. 121, 115–119.

Zhang, R., Han, P., Yang, H., Ouyang, K., Lee, D., Lin, Y.F., Ocorr, K., Kang, G., Chen, J., Stainier, D.Y., et al. (2013). In vivo cardiac reprogramming contributes to zebrafish heart regeneration. Nature 498, 497–501.

